# CASLFA: CRISPR/Cas9-mediated lateral flow nucleic acid assay

**DOI:** 10.1101/702209

**Authors:** Xusheng Wang, Erhu Xiong, Tian Tian, Meng Cheng, Wei Lin, Jian Sun, Xiaoming Zhou

**Affiliations:** College of Biophotonics & College of Life Science, South China Normal University, Guangzhou 510631, China; College of Veterinary Medicine, South China Agricultural University, Guangzhou 510642, China

**Keywords:** Lateral flow assay, CRISPR/Cas9, *Listeria monocytogenes* detection, GMOs detection, ASFV detection

## Abstract

The lateral flow assay is one of the oldest and most convenient analytical techniques for analyzing the immune response, but its applicability to precise genetic analyses is limited by the tedious and inefficient hybridization steps. Here, we have introduced a new version of the lateral flow assay, termed Cas9-mediated lateral flow nucleic acids assay (CASLFA), to address such issues. In this study, CASLFA is utilized to identify *Listeria monocytogenes*, genetically modified organisms (GMOs), and African swine fever virus (ASFV) at a sensitivity of hundreds of copies of genome samples with high specificity within 1 h. CASLFA satisfies some of the characteristics of a next-generation molecular diagnostics tool due to its rapidity and accuracy, allowing for point-of-care use without the need for technical expertise and complex ancillary equipment. This method has great potential for analyzing genes in resource-poor or nonlaboratory environments.

## Introduction

Due to its high sensitivity and excellent specificity, molecular diagnosis has shown great potential in applications in the fields of control and prevention of human and animal diseases, food safety detection, and environmental monitoring^1-3^. Some of these diagnostic techniques are playing key roles in clinical applications, such as prenatal testing^4^, oncogene mutation testing^5^, diagnosis of infectious diseases^6^, and identification of microorganisms^7^. Currently, gene sequencing, polymerase chain reaction (PCR), and *in situ* hybridization are the most widely used technologies. However, the dependence on expensive instruments and skilled technicians makes them unavailable in resource-limited laboratories or when rapid detection is needed. Lateral flow assays, such as lateral flow immunoassays, are one of the most convenient point-of-care testing (POCT) techniques and was launched commercially more than 30 years ago^8-14^. In recent years, lateral flow assays have been adopted for nucleic acids analyses that are rapidly being developed for gene identification^15, 16^. In a typical lateral flow nucleic acid assay (LFA), the analytes, such as double-stranded nucleic acids, are amplified by PCR or isothermal techniques using primers with two different tags, which are responsible for binding to their affinity ligand for capture on a paper substrate and signal output for detection^17, 18^. However, the major limitation of the existing LFAs is that the method fails to distinguish target-dependent from target-independent amplicons, such as primer dimers, thus leading to false-positive signals. A precise nucleic acid analysis has been achieved by designing a probe sequence for complementary hybridization^19^. Before lateral flow detection, the obtained amplicons are initially denatured at a high temperature to separate the strands and then renatured at a low temperature for probe-amplicon hybridization. However, tedious hybridization and manipulation steps and limited probe accessibility caused by the renaturation and formation of secondary structures make a conventional hybridization assay time-consuming and inefficient. Asymmetric nucleic acid amplification allows the production of single-stranded nucleic acids to be hybridized in a method compatible with mild temperature conditions, but at the expense of a reduced amplification efficiency^20^. Overall, the lack of convenience, rapidity, and efficiency has rendered the conventional LFA a non-user-friendly approach in practical applications. Thus, a novel LFA that addresses the problems mentioned above must be developed. This new approach will hopefully pave the way for the application of LFA technology to clinical or other molecular diagnostics platforms.

CRISPR/Cas technology has revolutionized various fields of molecular biology, such as gene editing, gene regulation, and gene imaging, and is showing great promise in the field of molecular diagnostics^21^. Currently, the application of CRISPR-Cas technology to genetic detection that relies on CRISPR/Cas12a, CRISPR/Cas13a, and CRISPR/Cas14a systems has been extensively developed^1, 3^. All of these methods depend on an indirect measurement of *trans*-cleavage activity triggered by target sequence recognition. However, *trans*-cleavage activity may be inhibited or nonspecifically activated by target-independent factors^22^.

The CRISPR/Cas9 system possesses excellent DNA recognition capabilities but does not possess target-dependent *trans*-cleavage activity; therefore, direct detection methods can be developed^23-25^. On the other hand, only CRISPR/Cas12a and CRISPR/Cas9 system are available for dsDNA recognition. The recognition of dsDNA by CRISPR/Cas12a depends on the protospacer adjacent motif (PAM) site, TTTN, adjacent to the target dsDNA^26-29^. This site is not detected as frequently in the genomic sequence as the PAM sequence of NGG or, to a lesser extent, NG for CRISPR/Cas9, which occurs once in every 8 bps or 4 bps of random DNA and thus is less restrictive than TTTN^30, 31^. We envisioned the use of CRISPR/Cas9 as a genetic recognition component to develop a more general genetic detection platform.

Using the CRISPR/Cas9 tool, our laboratory and other groups have developed new methods for genotyping pathogens and discriminating single nucleotide polymorphisms (SNPs) based on CRISPR/Cas9-mediated cleavage^32-35^. Very recently, two isothermal nucleic acid amplification methods based on the Cas9 nickase have also been developed and require a pair of Cas9/sgRNAs, two or three nucleases, accessory proteins, and expensive fluorescent probes^36, 37^. Successful detection of dsDNA has also been achieved by binding based on dead Cas9 (dCas9)^32-35^. Despite this continuous progress, the highly complex the biosensor construction process or cost of this technique hampers its widespread application, particularly at the point of care. In this study, we aim to develop a simplified nucleic acid assay that minimizes the reliance on instruments and professionals. This core of the technique is to integrate CRISPR/Cas9 recognition into a lateral flow detection platform; we thus termed this technology the Cas9-mediated lateral flow nucleic acid assay (CASLFA).

## Results and Discussion

Structure analyses have shown that PAM-dependent CRISPR/Cas9 recognition results in a sequential, stepwise unwinding, subsequently releasing the nontargeted strand of the target dsDNA as a single strand^38^. We thus hypothesized that the released nontargeted single strand triggered by CRISPR/Cas9 recognition can be used as the template for the probe (such as AuNP-DNA probes used in this study) hybridization. Based on this premise, we integrated the CRISPR/Cas9 system with a lateral flow device to develop a novel dsDNA detection technique (Figure 1A). We termed this strategy the DNA unwinding-based hybridization assay. In this strategy, the lateral flow device, which consists of a sample pad, a conjugate pad, test line, control line, and the absorbent pad, was designed (Figure 1A). An AuNP-DNA probe based on poly adenine (A)-Au affinity labeling was designed to coordinate with the DNA unwinding-based hybridization assay (Figure S1). The DNA sequence used to prepare the AuNP-DNA probe contains a labeling region, a control line hybridization region and a signal probe hybridization region, respectively (Figure 1A). In the DNA unwinding-based hybridization assay, genomic samples are amplified by PCR or isothermal reactions, such as recombinase polymerase amplification (RPA), using biotinylated primers. After a short incubation of the biotinylated amplicons with the designed Cas9/sgRNA for binding, the resulting cas9/sgRNA-biotinylated amplicons are then trickled onto the sample pad. Under capillary force, fluid flow is produced and the complexes initially flow laterally to the position of the conjugate pad, where they encounter and recognize the preassembled AuNP-DNA probes. In an automatic lateral flow detection format, Cas9/sgRNA-biotinylated amplicons-AuNP-DNA probe complexes continue to flow through the test line and are captured by the coated streptavidin. At the same time, excess AuNP-DNA probes flow through the lateral flow device and are trapped at the control line by hybridizing with precoated DNA probes. Color development and results are observed three minutes after sample application. As we anticipated, CASLFA works well for the EGFR gene used as an example, confirming the success of the DNA unwinding-based hybridization assay (Figure 1B).

**Figure 1.**
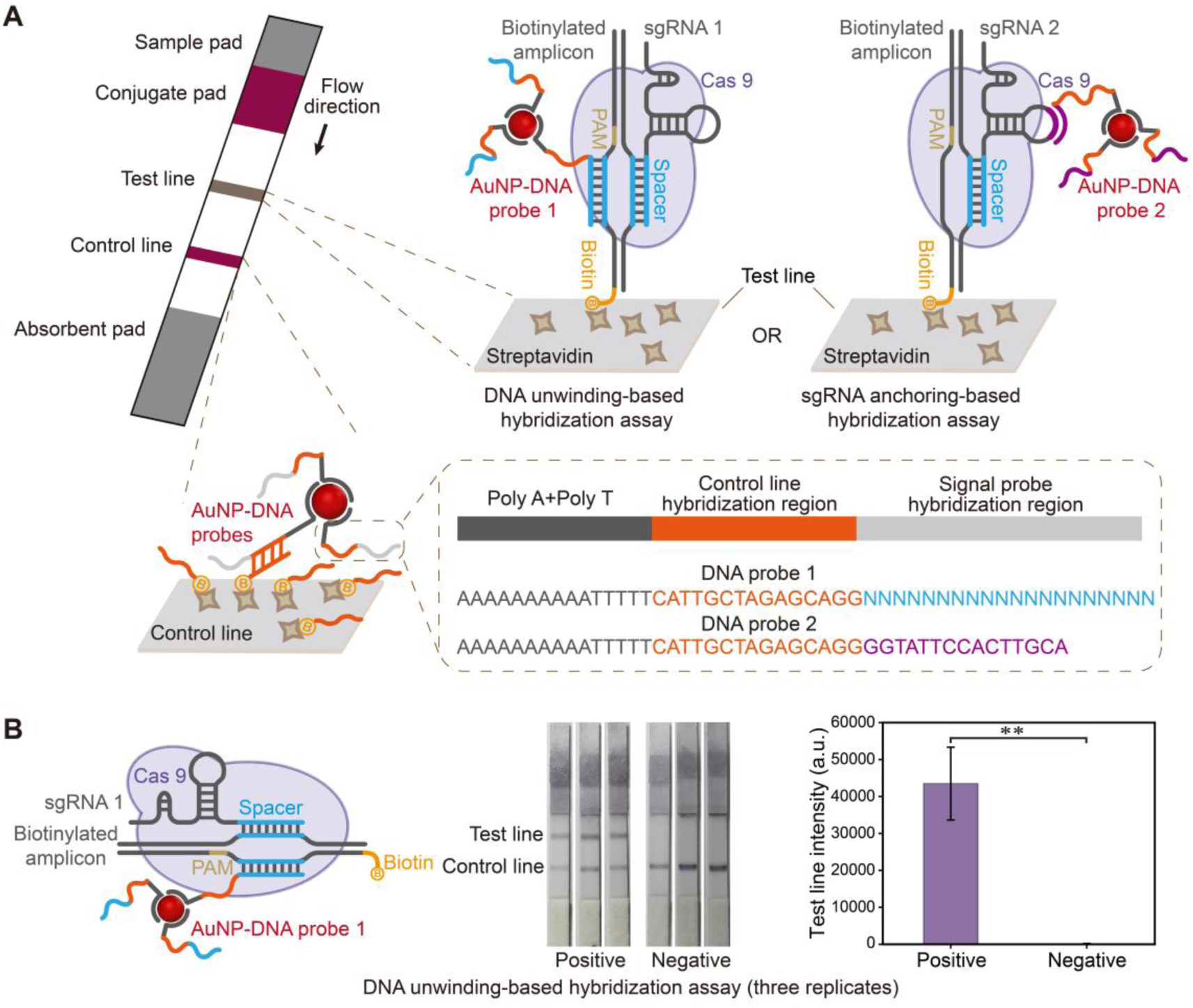
Schematic illustrating the developed CASLFA method. (A) Design of the lateral flow device. Left panel: The lateral flow device contains a sample pad, a conjugate pad, a test line, a control line, and an absorbent pad. The flow direction is represented by the arrow. AuNP-DNA probes were preassembled into the conjugate pad. Right panel: In the DNA unwinding-based hybridization assay, biotinylated amplicon-Cas9/sgRNA complexes were captured by the streptavidin-coated test line and AuNP-DNA probes were hybridized with the single-strand region of the amplicon released by Cas9/sgRNA-mediated unwinding. In the sgRNA anchoring-based hybridization assay, a sgRNA was designed to contain a universal sequence in the stem-loop region for AuNP-DNA probes hybridization. Biotinylated amplicon-Cas9/sgRNA complexes were captured by the streptavidin-coated test line and AuNP-DNA probes hybridized with the universal sequence in stem-loop region of the sgRNA. Lower panel: DNA probe 1 or 2 used to prepare AuNP-DNA contains three functional regions. Poly A + poly T: Poly A is used for affinity labeling with Au and poly T functions as the linker. The orange area represents the portion used for hybridization with the embedded probe in the Control line. The blue area (in DNA probe 1) and purple area (in DNA probe 2) represent the portions used for hybridization with the single-stranded region of the amplicon released by Cas9/sgRNA-mediated unwinding or the universal sequence in the stem-loop region of the sgRNA. (B) CASLFA method based on the DNA unwinding-based hybridization assay (left panel). Representative photographs taken from the CASLFA of EGFP amplicons (middle panel). CASLFA test line intensity from three independent experiments (right panel). (n = 3 technical replicates, two-tailed Student’s t-test; **, p < 0.01; bars represent means ± SEM)

Next, we planned to use the CASLFA method for transgene detection, pathogen identification, and virus detection. However, in the DNA unwinding-based hybridization assay, customized AuNP-DNA probes must be designed and labeled for different application purposes, potentially resulting in differences in detection performance caused by the variation in hybridization efficiency. In addition, the design, labeling, and assembly of AuNP-DNA probes in the lateral flow device also increase the time and cost of the CASLFA method. This limitation inspired us to develop a universal probe system in which equal detection efficiency will be maintained and the same lateral flow device can be used to detect different targets, thus endowing the CASLFA method with greater application prospects.

The early version of the S*treptococcus pyogenes* type II CRISPR system requires the nuclease Cas9, a targeting crRNA and an additional *trans*-activating crRNA to function. Subsequently, the crRNA and *trans*-activating crRNA were fused into a single guide RNA (sgRNA)^39^. The sgRNA contains constant scaffold sequences and variable recognition sequences. Numerous studies have also modified scaffold sequences to achieve better efficiency and diverse applications^40-42^. For example, the sgRNA scaffold has been engineered to contain an aptamer sequence that anchors to RNA binding proteins to regulate gene expression^43^. In the present study, we attempted to modify the scaffold sequence of the conventional sgRNA to obtain a universal hybridization site for AuNP-DNA probes. As shown in Figures 1A and 2A, we redesigned the sgRNA scaffold by inserting an additional sequence to generate an extended hairpin structure. We postulated that the extended hairpin structure would serve as an anchoring site for recruiting AuNP-DNA probes that had been preassembled into the lateral flow device, enabling the development of a universal sgRNA anchoring-based hybridization assay.

We engineered the sgRNA (sgRNA2) to contains a 15 bp extended hairpin structure as a probe for EGFP sequences to verify the feasibility of the sgRNA anchoring-based hybridization assay. An unmodified sgRNA, sgRNA1, was used as a reference for comparison. Based on the native polyacrylamide gel electrophoresis (PAGE) experiments, the two sgRNAs presented a remarkably different electrophoretic migration behavior, potentially due to the different lengths and conformations (Figure 2B). We then assembled Cas9 with the two sgRNAs to evaluate their cleavage efficiency. Cas9 is a single-turnover enzyme and the product is released at exceedingly slow rate after Cas9/sgRNA cleavage. Single-molecule studies have reported that separations can take up to several hours^44^. We thus analyzed the cleavage products using PAGE under denaturing and nondenaturing conditions. Both sgRNA1 and sgRNA2 worked well, while sgRNA2 appeared to be more efficient. An electrophoretic analysis of the cleavage products under nondenaturing conditions did not detect a characteristic cleaved band. However, the amplicon band disappeared, indicating that the cleaved products were still bound to the cas9/sgRNA complex, thus causing electrophoretic retention (Figure 2B). Since CASLFA relies on CRISPR/Cas9 recognition and binding rather than cleavage, we anticipate that both Cas9 and dCas9 can be adopted. As shown in Figure 2C, a similar detection performance was obtained when both Cas9-sgRNA2 and dCas9-sgRNA2 were used as the recognition modes, indicating that CASLFA is compatible with both Cas9 and dCas9 systems. Furthermore, we tested the Cas9-based CASLFA system under denaturing and nondenaturing conditions. The test line intensities were weaker after denaturation, confirming the adhesive properties of the cas9/sgRNA substrate after the cleavage reaction (Figure S2).

**Figure 2.**
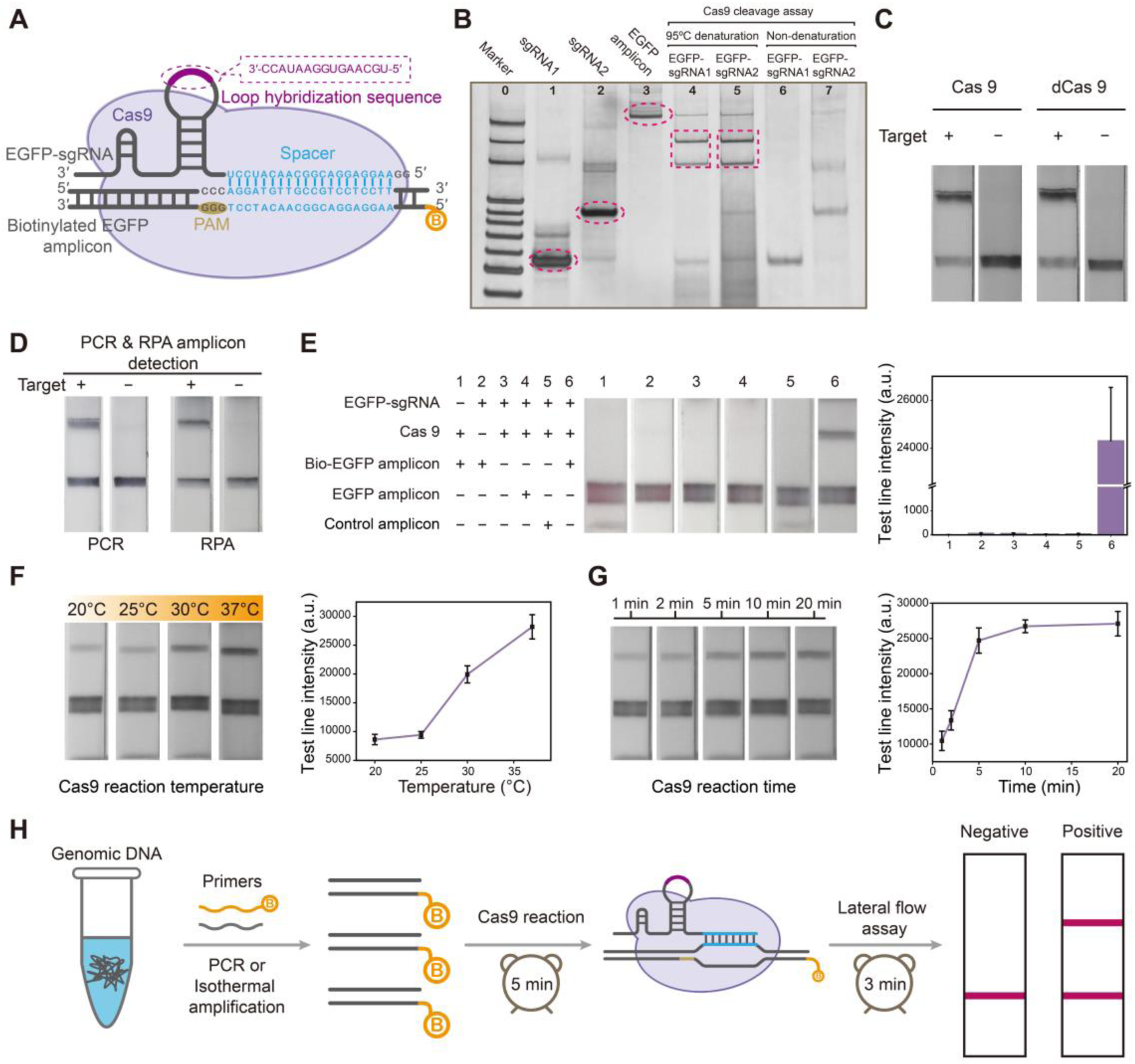
Verification of the universal probe strategy based on the sgRNA anchoring strategy. (A) EGFP amplicon recognition by the modified Cas9/gRNA system. (B) PAGE analysis of the conventional sgRNA and modified sgRNA, EGFP amplicon, and Cas9 cleavage assay. (C) CASLFA method based on Cas9 and dCas9. (D) CASLFA method for detecting PCR and RPA amplicons. (E) Component deletion experiments for corroborating the CASLFA concept. Right panel: Quantification of the test line intensity. (F) Evaluation of the temperature dependence of the Cas9 reaction. Right panel: Quantification of test line intensity. (G) Evaluation of the time dependence of the Cas9 reaction. Right panel: Quantification of test line intensity. (H) Workflow of the developed CASLFA method. (n = 3 technical replicates; bars represent mean ± SEM)

The CASLFA method was also further verified by testing its compatibility with PCR and RPA methods. We used biotinylated PCR or RPA primers to amplify the same target region and ensure that both PCR and RPA products were recognized by the same sgRNA. Both PCR and RPA were well adapted to the CASLFA technique (Figure 2D). Notably, PCR requires temperature cycling instruments and longer amplification times (approximately 1 hour), but requires cheaper reagents and exhibits better stability. RPA reagents are expensive, but less than 20 min for amplification and a simpler temperature control device (such as a metal bath) are required. Nevertheless, the CASLFA method combined with either PCR or RPA technologies can achieve diverse diagnostic needs.

Next, we performed component deletion experiments to further corroborate the CASLFA concept. The CASLFA method was only successful when the system contained Cas9, the correct sgRNA, and biotinylated target DNA (Figure 2E). We next evaluated two important parameters, Cas9-sgRNA reaction temperature and time, for the CASLFA method. Considering its future applications in nonlaboratory environments, we tested the performance of CASLFA at temperatures ranging from 20 to 37°C. Cas9-sgRNA works well at all temperature ranges, but displays the highest efficiency at 37°C (Figure 2F). On the other hand, Cas9-sgRNA quickly searches for its target DNA. We thus analyzed incubation times ranging from 1 to 20 min, and 5 minutes was sufficient to observe a saturated detection signal (Figure 2G).

After conceptual verification, we summarized the detection procedure of CASLFA method (Figure 2H). Following gene amplification (20 min to 1 hour. depending on the amplification method), the test sample was incubation with Cas9-sgRNA for a short period (5 min) and then transferred to the lateral flow device. CASLFA then proceeds automatically under capillary flow, which requires approximately 3 min to complete. The results can be analyzed using the naked eye. If the RPA amplification method is used, the whole CASLFA process can be completed in 1 hour at a constant temperature (37°C).

After developing a universal CASLFA system, we next evaluated the detection performance of the CASLFA method. We selected *Listeria monocytogenes* (a common and high-risk food-borne pathogen), transgenic 35S promoter (a most commonly used target for GMO detection), and African swine fever virus (ASFV, a deadly pig virus outbreak worldwide starting in 2018) to determine the versatility and sensitivity of CASLFA method. For the detection of *Listeria monocytogenes*, the *hlyA* gene was used as a target sequence and the corresponding CRISPR/Cas9 system was designed (Figure 3A). Genomic samples from *Listeria monocytogenes* were diluted 10-fold in a gradient and then amplified by PCR. The amplified product was subjected to the CASLFA method and the results were visually observed from the lateral flow device (Figure 3B). As shown in Figure 3C, the detection limit of the naked eye is as low as approximately 150 copies of the genome sample without a optimizing amplification parameters. At the same time, the CASLFA for *Listeria monocytogenes* detection was also further verified in combination with the RPA method and achieved a similar sensitivity to PCR (Figure S3). Subsequently, the specificity of the CASLFA method was also evaluated. As shown in Figure 3D, detection results visually the excellent specificity of the developed CASLFA method for *Listeria monocytogenes* among five other interfering bacteria.

**Figure 3.**
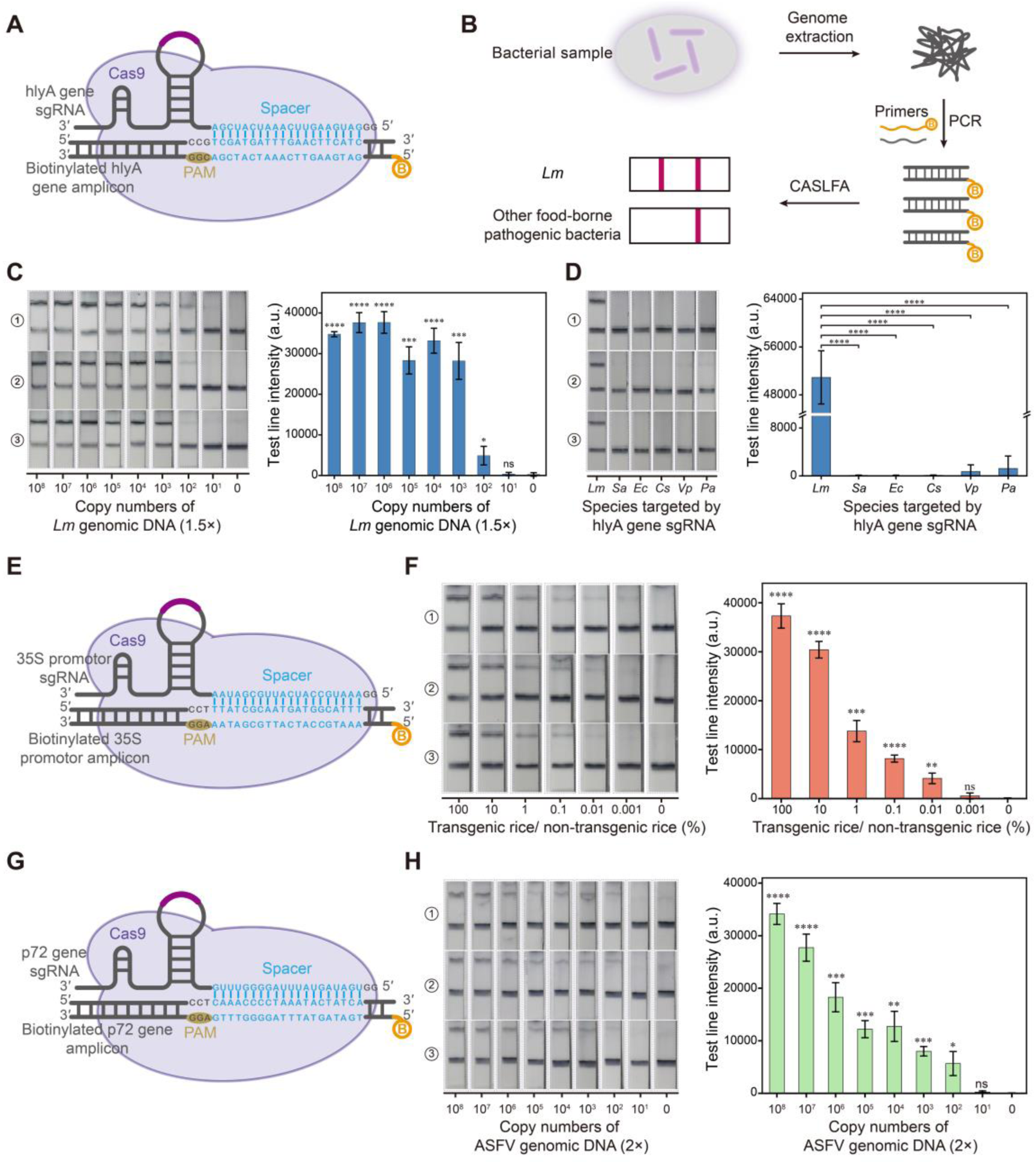
Applications and the analytical performance of the developed CASLFA method. (A) Using the CASLFA method, the *Listeria monocytogenes hlyA* gene was recognized by the designed Cas9/sgRNA system. (B) Workflow of *Lm* detection using the CASLFA method. *Lm, Listeria monocytogenes*. (C) Evaluation of the sensitivity of the CASLFA method for *Lm* detection. Right panel: Quantification of the test line intensity. (D) Evaluation of the specificity of the CASLFA method for *Lm* detection. Right panel: Quantification of test line intensity. *Sa, Staphylococcus aureus*; *Ec, Escherichia coli*; *Cs, Cronobacter sakazakii*; *Vp, Vibrio parahaemolyticus*; *Pa, Pseudomonas aeruginosa*. (E) Use of the CASLFA method to detect GMOs. The 35S promotor sequence was recognized by the designed Cas9/sgRNA system. (F) Evaluation of the sensitivity of the CASLFA method for GMO detection. Left panel: Photograph showing strip detection results in the presence of various percentages of transgenic rice DNA. Right panel: Quantitation of test line intensity from the strip detection. (G). Use of the CASLFA method for the detection of ASFV. The P72 gene sequence was recognized by the designed Cas9/sgRNA system. (H). Evaluation of the sensitivity of the CASLFA method for ASFV detection. Left panel: Photograph showing strip detection results in the presence of various copy numbers of ASFV DNA. Right panel: Quantitation of test line intensity from the strip detection. (n = 3 technical replicates, two-tailed Student’s t-test; ns, not significant; *, p < 0.05; **, p < 0.01; ***, p < 0.001; and ****, p < 0.0001; bars represent means ± SEM)

Next, we evaluated the ability of the CASLFA system to detect the 35S sequence of transgenic rice. We mixed transgenic rice with nontransgenic rice at different ratios (from 0.001% to 100%) and milled the rice into a powder for nucleic acid extraction and PCR. Based on observations with the naked eye and the grayscale analysis, 0.01% of transgenic samples were easily detectable. The sensitivity of the method is appropriate for the detection of GMO components ranging from 0.1% to 1% worldwide. Finally, various copy numbers of genomic DNA samples from ASFV samples were also subjected to the CASLFA analysis. Sensitivity was evaluated by both observations with the naked eye and a grayscale analysis, and a limit of detection of 200 copies of viral genomic DNA was obtained.

## Conclusion

In summary, CASLFA represents a simple and rapid method for the detection of genetic targets using the naked eye. Using a pathogenic microorganism, GMOs, and virus as examples, CASLFA achieves sensitivity levels as low as hundreds of gene copies, which is comparable to most PCR-based techniques. Importantly, by employing the sgRNA anchoring-based hybridization assay, the AuNP-DNA probe is universal, and the lateral flow device preassembled with the AuNP-DNA probe can be applied to detect any target gene. After nucleic acid extraction, the CASLFA method for gene detection requires half an hour (in combination with RPA) to approximately one hour (in combination with PCR) to complete. Although this study does not cover the extraction of nucleic acids, recent advances indicated that nucleic acid preparation can be completed in a few minutes in nonlaboratory environments^45, 46^. Therefore, with combined with these nucleic acid extraction strategies, we envisage that CASLFA will be able to complete the entire genetic detection process in one hour, which meet the needs for rapid genetic detection in most cases. Other significant merits of CASLFA include no need of a separation process and no temperature ramping procedure during the detection step. Thus, CASLFA does not require any complex equipment, making this approach particularly suitable for POC testing. Thus, we concluded that the CASLFA method satisfies some of the characteristics of next-generation molecular diagnostics because it is inexpensive, accurate, and rapidly yields results, allowing for point-of-care use without the need for technical expertise and ancillary equipment and confirming its great potential for functional gene analyses.

## Experimental Section

Experimental details are provided in the Supporting Information.

## Acknowledgment

This work was supported by the National Natural Science Foundation of China (Grant 21475048; 21874049), the National Science Fund for Distinguished Young Scholars of Guangdong Province (Grant 2014A030306008), and the Special Support Program of Guangdong Province (Grant 2014TQ01R599).

## References

1. Li, Y.; Li, S.; Wang, J.; Liu, G. Trends in Biotechnology 2019, 37, (7), 730–743.

2. Korf, B. R.; Rehm, H. L. Jama 2013, 309, (14), 1511–1521.

3. Chertow, D. S. Science 2018, 360, (6387), 381–382.

4. Nicolaides, K. H.; Syngelaki, A.; Ashoor, G.; Birdir, C.; Touzet, G. American Journal of Obstetrics and Gynecology 2012, 207, (5), 374. e1-374. e6.

5. Koboldt, D. C.; Zhang, Q.; Larson, D. E.; Shen, D.; McLellan, M. D.; Lin, L.; Miller, C. A.; Mardis, E. R.; Ding, L.; Wilson, R. K. Genome Research 2012, 22, (3), 568–576.

6. Lee, W. G.; Kim, Y.-G.; Chung, B. G.; Demirci, U.; Khademhosseini, A. Advanced Drug Delivery Reviews 2010, 62, (4-5), 449–457.

7. Chung, H. J.; Castro, C. M.; Im, H.; Lee, H.; Weissleder, R. Nature Nanotechnology 2013, 8, (5), 369–375.

8. Bahadır, E. B.; Sezgintürk, M. K. TrAC Trends in Analytical Chemistry 2016, 82, 286–306.

9. Gao, Z.; Ye, H.; Tang, D.; Tao, J.; Habibi, S.; Minerick, A.; Tang, D.; Xia, X. Nano Letters 2017, 17, (9), 5572–5579.

10. Liu, J.; Mazumdar, D.; Lu, Y. Angewandte Chemie International Edition 2006, 45, (47), 7955–7959.

11. Qin, Z.; Chan, W. C.; Boulware, D. R.; Akkin, T.; Butler, E. K.; Bischof, J. C. Angewandte Chemie International Edition 2012, 51, (18), 4358–4361.

12. Reboud, J.; Xu, G.; Garrett, A.; Adriko, M.; Yang, Z.; Tukahebwa, E. M.; Rowell, C.; Cooper, J. M. Proceedings of the National Academy of Sciences 2019, 116, (11), 4834–4842.

13. Tran, V.; Walkenfort, B.; König, M.; Salehi, M.; Schlücker, S. Angewandte Chemie International Edition 2019, 58, (2), 442–446.

14. Zhan, L.; Guo, S.-z.; Song, F.; Gong, Y.; Xu, F.; Boulware, D. R.; McAlpine, M. C.; Chan, W. C.; Bischof, J. C. Nano Letters 2017, 17, (12), 7207–7212.

15. Corstjens, P.; Zuiderwijk, M.; Brink, A.; Li, S.; Feindt, H.; Niedbala, R. S.; Tanke, H. Clinical Chemistry 2001, 47, (10), 1885–1893.

16. Mao, X.; Ma, Y.; Zhang, A.; Zhang, L.; Zeng, L.; Liu, G. Analytical Chemistry 2009, 81, (4), 1660–1668.

17. Crannell, Z.; Castellanos-Gonzalez, A.; Nair, G.; Mejia, R.; White, A. C.; Richards-Kortum, R. Analytical Chemistry 2016, 88, (3), 1610–1616.

18. Li, S.; Gu, Y.; Lyu, Y.; Jiang, Y.; Liu, P. Analytical Chemistry 2017, 89, (22), 12137–12144.

19. Xu, Y.; Wei, Y.; Cheng, N.; Huang, K.; Wang, W.; Zhang, L.; Xu, W.; Luo, Y. Analytical Chemistry 2017, 90, (1), 708–715.

20. Xu, Y.; Liu, Y.; Wu, Y.; Xia, X.; Liao, Y.; Li, Q. Analytical Chemistry 2014, 86, (12), 5611–5614.

21. Pickar-Oliver, A.; Gersbach, C. A. Nature Reviews Molecular Cell Biology 2019, doi: 10.1038/s41580-019-0131-5

22. Tambe, A.; East-Seletsky, A.; Knott, G. J.; Doudna, J. A.; O’Connell, M. R. Cell Reports 2018, 24, (4), 1025–1036.

23. Hajian, R.; Balderston, S.; Tran, T.; deBoer, T.; Etienne, J.; Sandhu, M.; Wauford, N. A.; Chung, J.-Y.; Nokes, J.; Athaiya, M. Nature Biomedical Engineering 2019, 3, (6), 427–437.

24. Lee, H.; Choi, J.; Jeong, E.; Baek, S.; Kim, H. C.; Chae, J.-H.; Koh, Y.; Seo, S. W.; Kim, J.-S.; Kim, S. J. Nano Letters 2018, 18, (12), 7642–7650.

25. Yang, W.; Restrepo-Pérez, L.; Bengtson, M.; Heerema, S. J.; Birnie, A.; van der Torre, J.; Dekker, C. Nano Letters 2018, 18, (10), 6469–6474.

26. Chen, J. S.; Ma, E.; Harrington, L. B.; Da Costa, M.; Tian, X.; Palefsky, J. M.; Doudna, J. A. Science 2018, 360, (6387), 436–439.

27. Gootenberg, J. S.; Abudayyeh, O. O.; Kellner, M. J.; Joung, J.; Collins, J. J.; Zhang, F. Science 2018, 360, (6387), 439–444.

28. Li, S.-Y.; Cheng, Q.-X.; Wang, J.-M.; Li, X.-Y.; Zhang, Z.-L.; Gao, S.; Cao, R.-B.; Zhao, G.-P.; Wang, J. Cell Discovery 2018, 4, (1), 20.

29. Yan, W. X.; Hunnewell, P.; Alfonse, L. E.; Carte, J. M.; Keston-Smith, E.; Sothiselvam, S.; Garrity, A. J.; Chong, S.; Makarova, K. S.; Koonin, E. V. Science 2019, 363, (6422), 88–91.

30. Hu, J. H.; Miller, S. M.; Geurts, M. H.; Tang, W.; Chen, L.; Sun, N.; Zeina, C. M.; Gao, X.; Rees, H. A.; Lin, Z. Nature 2018, 556, (7699), 57–63.

31. Liu, J.-J.; Orlova, N.; Oakes, B. L.; Ma, E.; Spinner, H. B.; Baney, K. L.; Chuck, J.; Tan, D.; Knott, G. J.; Harrington, L. B. Nature 2019, 566, (7743), 218–223.

32. Gu, W.; Crawford, E. D.; O’Donovan, B.; Wilson, M. R.; Chow, E. D.; Retallack, H.; DeRisi, J. L. Genome Biology 2016, 17, (1), 41.

33. Huang, M.; Zhou, X.; Wang, H.; Xing, D. Analytical Chemistry 2018, 90, (3), 2193–2200.

34. Lee, S. H.; Yu, J.; Hwang, G. H.; Kim, S.; Kim, H.; Ye, S.; Kim, K.; Park, J.; Park, D.; Cho, Y. Oncogene 2017, 36, (49), 6823–6829.

35. Quan, J.; Langelier, C.; Kuchta, A.; Batson, J.; Teyssier, N.; Lyden, A.; Caldera, S.; McGeever, A.; Dimitrov, B.; King, R. Nucleic Acids Research 2019, doi: 10.1093/nar/gkz418.

36. Wang, T.; Liu, Y.; Sun, H. H.; Yin, B. C.; Ye, B. C. Angewandte Chemie International Edition 2019, 58, (16), 5382–5386.

37. Zhou, W.; Hu, L.; Ying, L.; Zhao, Z.; Chu, P. K.; Yu, X.-F. Nature Communications 2018, 9, (1), 5012.

38. Jiang, F.; Zhou, K.; Ma, L.; Gressel, S.; Doudna, J. A. Science 2015, 348, (6242), 1477–1481.

39. Jinek, M.; Chylinski, K.; Fonfara, I.; Hauer, M.; Doudna, J. A.; Charpentier, E. Science 2012, 337, (6096), 816–821.

40. Jusiak, B.; Cleto, S.; Perez-Pinera, P.; Lu, T. K. Trends in Biotechnology 2016, 34, (7), 535–547.

41. Shao, S.; Zhang, W.; Hu, H.; Xue, B.; Qin, J.; Sun, C.; Sun, Y.; Wei, W.; Sun, Y. Nucleic Acids Research 2016, 44, (9), e86–e86.

42. Zhang, K.; Deng, R.; Teng, X.; Li, Y.; Sun, Y.; Ren, X.; Li, J. Journal of the American Chemical Society 2018, 140, (36), 11293–11301.

43. Zalatan, J. G.; Lee, M. E.; Almeida, R.; Gilbert, L. A.; Whitehead, E. H.; La Russa, M.; Tsai, J. C.; Weissman, J. S.; Dueber, J. E.; Qi, L. S. Cell 2015, 160, (1-2), 339–350.

44. Ma, H.; Tu, L.-C.; Naseri, A.; Huisman, M.; Zhang, S.; Grunwald, D.; Pederson, T. Journal of Cell Biology 2016, 214, (5), 529–537.

45. Zou, Y.; Mason, M. G.; Wang, Y.; Wee, E.; Turni, C.; Blackall, P. J.; Trau, M.; Botella, J. R. PLoS biology 2017, 15, (11), e2003916.

46. Paul, R.; Saville, A. C.; Hansel, J. C.; Ye, Y.; Ball, C.; Williams, A.; Chang, X.; Chen, G.; Gu, Z.; Ristaino, J. B. ACS Nano 2019, 13, (6), 6540–6549.

