## Supporting Information for "CASLFA: CRISPR/Cas9-mediated lateral flow nucleic acid assay"

Materials

The nitrocellulose membrane was purchased from Sartorius. The sample pad, bonding pad, bottom plate and absorbent pad were purchased from Shanghai Jieyi Biotechnology Co., Ltd. (Shanghai, China). Bovine serum albumin (BSA) was purchased from Sigma-Aldrich (St. Louis, MO, USA). Twist Amp® Basic Kit used for RPA reaction was purchased from TwistDx (England). Streptavidin, sucrose, sodium phosphate, Tween-20, Triton X-100, Tris-HCl, 20x saline sodium citrate (SSC) buffer, 20x phosphate buffer saline (PBS), and NaCl were purchased from Sangon Biotech (Shanghai, China). Disodium hydrogen phosphate, sodium dihydrogen phosphate, concentrated nitric acid, and hydrochloric acid were purchased from Guangzhou Chemical Reagent Factory (Guangzhou, China). The T7 RNA polymerase, T7 reaction buffer, and NTPs required for RNA transcription were purchased from Bio-Lifesci (Guangzhou, China). Premix Taq™ (TaKaRa Taq™ Version 2.0) and PrimeSTAR® Max DNA Polymerase used for PCR were purchased from Takara (Japan). All oligonucleotide sequences listed in Table S1 were synthesized by Sangon Biotech (Shanghai, China). The reagents used for protein expression and purification were purchased from Abiotech (Jinan, China). The pET28a/Cas9-Cys plasmid used to prepare S. pyogenes Cas9 was a gift from Hyongbum Kim (Addgene plasmid # 53261, http://n2t.net/addgene:53261, RRID: Addgene_53261). The pET302-6His-dCas9-Halo plasmid used to prepare dCas9 was a gift from Timothée Lionnet (Addgene plasmid #72269, http://n2t.net/addgene:72269, RRID: Addgene_72269). Instruments used in high-speed centrifugation and PCR experiments were purchased from Eppendorf. The XYZ three-dimensional spray gold instrument was purchased from Shanghai Jinbiao Biotechnology Co., Ltd. (Shanghai, China). The agarose and PAGE electrophoresis experiments were performed using equipment from Beijing Liuyi Instruments (Beijing, China), and the instrument used for imaging was a BG-gdsAUTO320 gel imaging analysis system (Beijing, China). The instrument used for the absorption spectroscopy experiments was a SpectraMax iD5 Multi-Mode Microplate Reader (Molecular Devices, USA). All other buffers were prepared using ultrapure water (> 18.25 MΩ/cm).

**Experimental Procedures**

**Preparation of AuNPs**

AuNPs with a diameter of 13 nm were prepared using the sodium citrate reduction method. The glassware required for the preparation process was thoroughly cleaned with aqua regia, then thoroughly rinsed with deionized water and dried. The preparation process is briefly described below. Ninety-eight milliliters of deionized water were into a flask containing a magnetic stir bar, 2 mL of a 50 mM HAuCl_4_ solution were added for a final HAuCl_4_ concentration of 1 mM, and then the mixture was heated on a heating type magnetic stirrer. After the solution boils, 10 mL of a 38.8 mM sodium citrate solution were quickly added. The color should change from light yellow to deep red. After heating for approximately 20 min, the solution is cooled to room temperature with stirring. Thus, AuNPs with a diameter of 13 nm were obtained (Figure S1).

**Preparation of AuNP-DNA probes**

The AuNP-DNA probes were prepared using the freezing method. Briefly, 500 μL of 13 nm AuNPs were added to 1 OD poly A-DNA probes and frozen at – 20°C for 1 hour. After thawing, the solution was centrifuged at 13000 rpm/min for 30 min at 4°C. The process was repeated 4 times. The buffer used to resuspend the pellet in the first three centrifugation steps was 0.1 M NaCl + 0.01 M PB buffer, and the pellet was resuspended in three volumes of distilled water and then centrifuged for the last step. At this time, the supernatant should be transparent. The supernatant was carefully aspirated and then the pellet was resuspended with 300 μL of buffer (20 mM Na_3_PO_4_, 5% BSA, 0.25% Tween-20, and 10% sucrose). The resulting AuNP-DNA probes were stored at 4°C in the dark until use (Figure S1).

**Cas9 and dCas9 protein expression and purification**

The Cas9 protein was expressed in *E. coli* Rosetta2 (DE3) cells cultured in Terrific Broth (TB) containing 34 μg/mL chloramphenicol and 50 μg/mL kanamycin. The cells were cultured overnight at 18°C until an OD_600_ = 0.6 was observed, after which 0.1 mM isopropyl-1-thio-b-D-galactopyranoside (IPTG) was added and cells were cultured at 26°C for 4 h [to induce protein expression](http://dict.youdao.com/w/inducible%20expression/" \l "keyfrom=E2Ctranslation). The collected bacterial solution was centrifuged to obtain a precipitate, and the precipitate was ultrasonically lysed with lysis buffer (20 mM Tris-HCl, pH 7.5, 1 M NaCl, and 10% glycerol). After centrifugation at 4°C, the supernatant purified on a nickel column (Abiotech, Jinan, China), washed with a washing solution (20 mM Tris-HCl, pH 7.5, 150 mM NaCl, 20 mM imidazole, and 10% glycerol) to remove the heterologous proteins, and eluted with elution buffer (20 mM Tris-HCl, pH 7.5, 150 mM NaCl, 500 mM imidazole, and 10% glycerol). The eluted Cas9 proteins were collected, dialyzed against dialysis solution (20 mM Tris-HCl, pH 7.5, 150 mM NaCl, and 50% glycerol), and stored at – 20°C. The dCas9 protein was expressed in *E. coli* Rosetta2 (DE3) cells cultured in TB containing 34 μg/mL chloramphenicol and 50 μg/mL ampicillin. The cells were cultured overnight at 18°C until an OD_600_ = 0.6 was observed, after which IPTG (0.1 mM) was added and incubated with the culture at 26°C for 4 h [to induce protein expression](http://dict.youdao.com/w/inducible%20expression/" \l "keyfrom=E2Ctranslation). Subsequent purification conditions and methods are identical to the Cas9 protein described above. The characterization of the purified Cas9 and dCas9 proteins is shown in Figure S4.

**Synthesis of sgRNA**

All sgRNAs described in the present study were transcribed *in vitro* from DNA templates using T7 RNA polymerase. The DNA template was obtained by a bridge-fill-in PCR method where the forward primer comprises a T7 promoter sequence and a 20 nt targeting sequence; the backward primers consist primarily of sequences encoding the 3' end of sgRNA scaffold. PCR was performed using PrimeSTAR® Max DNA Polymerase (Takara) under the following thermal cycling program: 95°C for 2 min, followed by 35 cycles of amplification at 95°C for 20 s, 63°C for 10 s, and 72°C for 45 s, a final extension at 72°C for 10 min and a 10°C hold. The PCR product served as a template for T7 RNA polymerase-mediated transcription. The transcription reaction was incubated at 37°C for 4 h. The [obtained](http://dict.youdao.com/w/obtained/" \l "keyfrom=E2Ctranslation) sgRNA was used immediately or stored at – 80°C. The characterization of the synthesized sgRNAs is shown in Figure S5.

**Preparation of the lateral flow device**

The lateral flow device consists of a sample pad, a bonding pad, a nitrocellulose membrane, an absorbent pad, and a bottom plate. The sample pad was composed of glass fibers, thoroughly wetted with a buffer (pH 8.0) containing 0.25% Triton X-100, 0.05 M Tris-HCl, and 0.15 M NaCl, and then dried at 37°C for 2 h. The material used for the bonding pad was also glass fibers that had been completely embedded with the AuNP-DNA probes and dried at 37°C for 2 h. The test line on the nitrocellulose membrane was sprayed with streptavidin, and the control line was sprayed with a streptavidin-biotinylated DNA probe solution. The distance between the test line and the control line was 7 mm, and then the nitrocellulose membrane (NC) film was dried at 37°C for 1 h. Finally, the sample pad, the bonding pad, the NC film and the absorption pad were attached to the bottom plate, each overlapping by 2 mm, and finally the lateral flow device with a width of 4 mm was cut and stored at 4°C.

**Genomic DNA extraction**

All bacterial genomic DNA used in the present study was extracted using the TIANamp Bacteria DNA kit (Beijing, China). The extracted DNA was stored at – 20°C until use. The DNA was extracted from two rice lines using a plant genomic DNA extraction kit (TianGen Plant Genomic DNA Kit, DP305). When extracting different proportions of transgenic rice, transgenic rice and nontransgenic rice were mixed in different mass ratios (for example, 1% transgenic rice samples are mixed with 1 g of transgenic rice and 99 g of nontransgenic rice), and then mixed samples were used for genome extraction. The extracted DNA was stored at – 20°C until use. Viral DNA templates from African swine fever virus were provided by Guangdong Provincial Centers for Disease Control and Prevention of China. These clinical samples were used as the source of the viral DNA templates and were collected from Huangpu District, Guangzhou City, Guangdong Province.

**Preparation of biotinylated amplicons**

(1) PCR amplification of the EGFP target DNA

The EGFP target fragment was obtained by PCR amplification using the pEGFP-N1 plasmid as a template and the primers shown in Table S1. The 50 μL PCR system contained 300 nM primer pairs, 25 μL of Premix Taq, 1 μL of the DNA template and deionized water. The PCR thermal cycling program was 95°C for 5 min, followed by 35 cycles of amplification at 95°C for 30 s, 60°C for 30 s, and 72°C for 1 min, a final extension at 72°C for 10 min and a 10°C hold. The amplified product was stored at – 20°C until use.

(2) PCR amplification of the *hlyA* gene from *Listeria monocytogenes*

The PCR primers designed to amplify the *Listeria* *monocytogenes hIyA* conserved gene (GenBank: KJ504109.1) are shown in Table S1. The 50 μL PCR system included 300 nM primer pairs, 25 μL of Premix Taq, 1 μL of different concentrations of the DNA template and deionized water. The PCR thermal cycling program was 95°C for 5 min, followed by 35 cycles of amplification at 95°C for 10 s, 51°C for 30 s, and 72°C for 45 s, a final extension at 72°C for 10 min and a 10°C hold. The amplified product was stored at – 20°C until use. The electrophoretic characterization of PCR products obtained from the amplification of different copy numbers of *Listeria monocytogenes* genomic samples is shown in Figure S6A. The electrophoretic characterization of the specificity of the amplification of the *hlyA* gene is shown in Figure S6B.

(3) PCR amplification of the 35S promoter of transgenic rice

PCR primers for the 35S promoter of transgenic rice are shown in Table S1. The 50 μL PCR system included 300 nM primer pairs, 25 μL of Premix Taq, 1 μL of different concentrations of DNA templates and deionized water. The PCR thermal cycling program was 95°C for 5 min, followed by 35 cycles of amplification at 95°C for 30 s, 51°C for 30 s, and 72°C for 45 s, a final extension at 72°C for 10 min and a 10°C hold. The amplified product was stored at – 20°C until use. The electrophoretic characterization of PCR products obtained from amplifying various percentages of transgenic rice DNA is shown in Figure S6C.

(4) PCR amplification of the African swine fever virus VP72 gene

PCR primers for the conserved VP72 gene of the African swine fever virus are shown in Table S1. The 50 μL PCR system included 300 nM primer pairs, 25 μL of Premix Taq, 1 μL of different concentrations of the DNA template and deionized water. The PCR thermal cycling program was 95°C for 15 min, followed by 40 cycles of amplification at 95°C for 10 s, 51°C for 30 s, and 72°C for 45 s, a final extension at 72°C for 10 min and a 10°C hold. The amplified product was stored at – 20°C until use. The electrophoretic characterization of PCR products obtained from amplifying various copy numbers of ASFV genomic samples is shown in Figure S6D.

(5) Recombinase polymerase amplification (RPA) of genomic DNA samples

RPA reactions were performed using a Twist Amp® Basic Kit (TwistDx). Briefly, the 50 µL RPA reaction contained 480 nM primer pairs, 29.5 µL of rehydration buffer, one freeze-dried reaction pellet, 1 µL of DNA template, and 2.5 µL of 280 mM MgAc (added last). RPA reactions were incubated at 37°C for 20 min. The electrophoretic characterization of RPA products obtained from amplifying varied copies of *Listeria monocytogenes* genomic samples is shown in Figure S3A.

***In vitro* CRISPR/Cas9 cleavage experiments**

Cas9 and sgRNA (1:1, 100 nM) were preincubated at 25°C for 10 min in reaction buffer (5 mM MgCl_2_) in a total volume of 8 µL. Afterwards, 2 µL of the EGFP amplicon (1 nM final concentration) were added to the mixture and incubated at 37°C for 1 h. Reactions were stopped by heating the samples to 95°C for 5 min. The cleavage products were analyzed using PAGE.

**CASLFA detection process**

The detection process includes the Cas9 reaction and a lateral flow detection process. First, 20 μL of the Cas9 reaction containing 200 nM Cas9, 150 nM sgRNA, 5 mM MgCl2 and 4 μL of biotinylated amplicons were incubated at 37°C for 5 min to obtain the Cas9-sgRNA-biotinylated amplicon complex. Thirty microliters of running buffer (components: 4x SSC, 0.05% Tween-20 (v/v), 1x PBS, and 1% BSA fraction 5) were added to the 20 μL Cas9 reaction solution, and then this mixture was to the sample pad of the lateral flow device in a dropwise manner. Subsequently, 50 μL of running buffer were added and the bands in the test line and control line appeared after 2 min. After completing the test, a photograph of the lateral flow device was captured, and the signal intensity of the test line was quantified using ImageJ software.

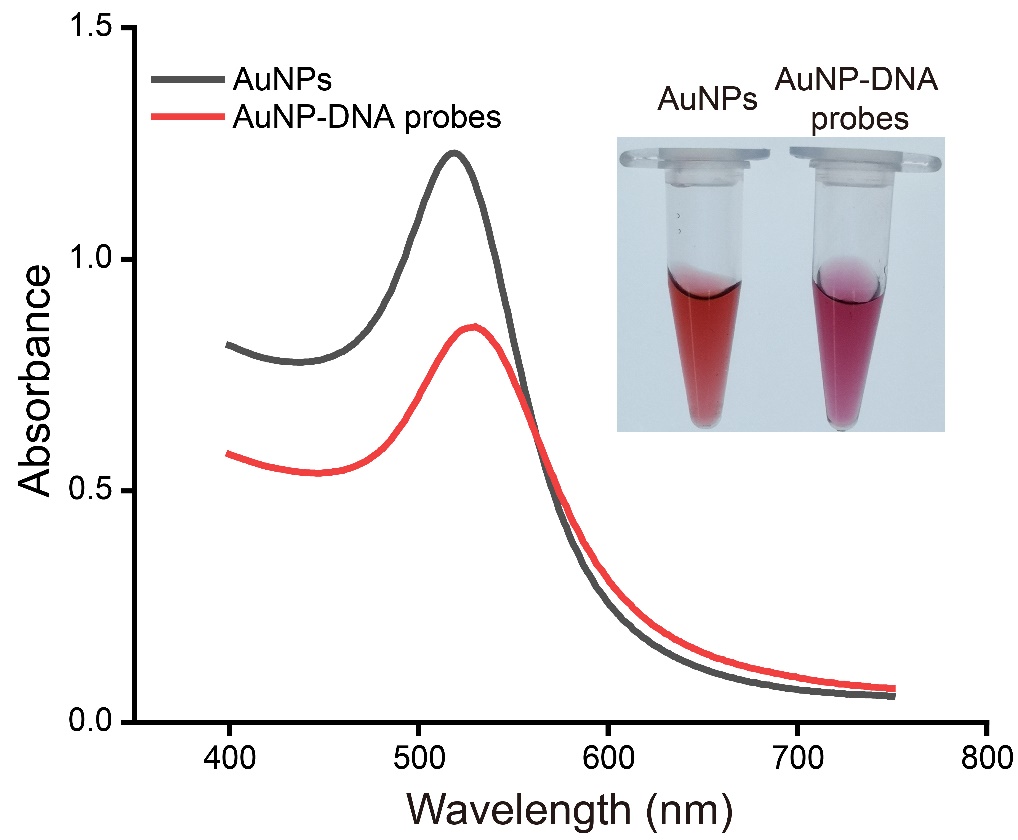

**Figure S1.** UV–Vis spectra and images (inset) showing the characterization of synthesized AuNPs and AuNP-DNA probes solution.

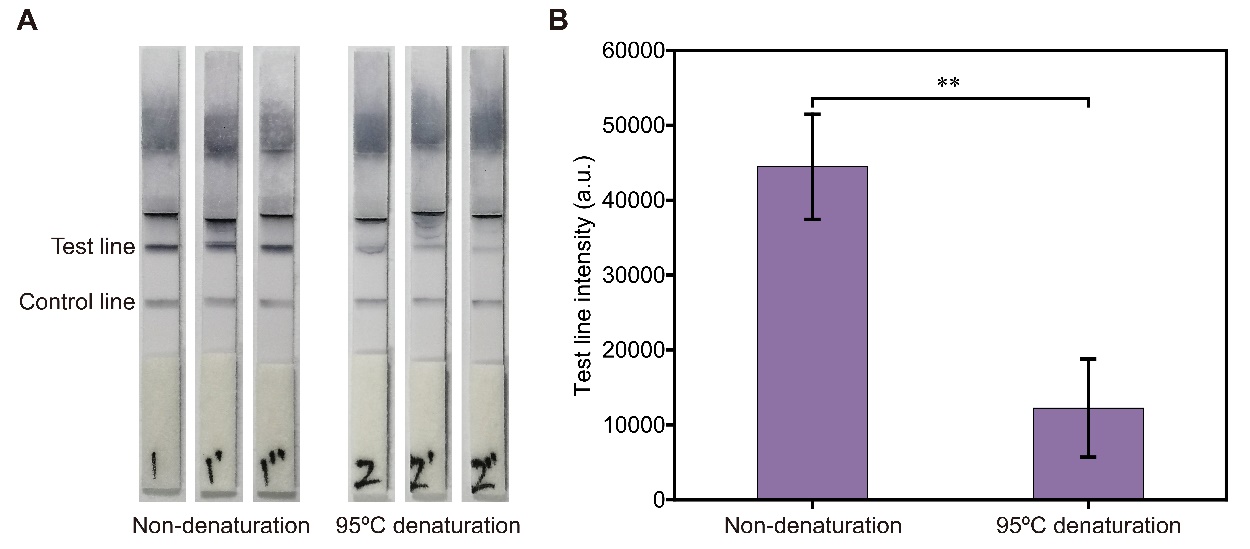

**Figure S2.** **Verification of the use of Cas9 for CASLFA under the denaturing and non-denaturing conditions.** (A) Photographs of lateral flow detection under nondenaturing and 95°C denaturing conditions (3 replicates). (B) Quantitation of test line intensities. (n = 3 technical replicates, two-tailed Student’s t-test; **, p < 0.01; bars represent means ± SEM)

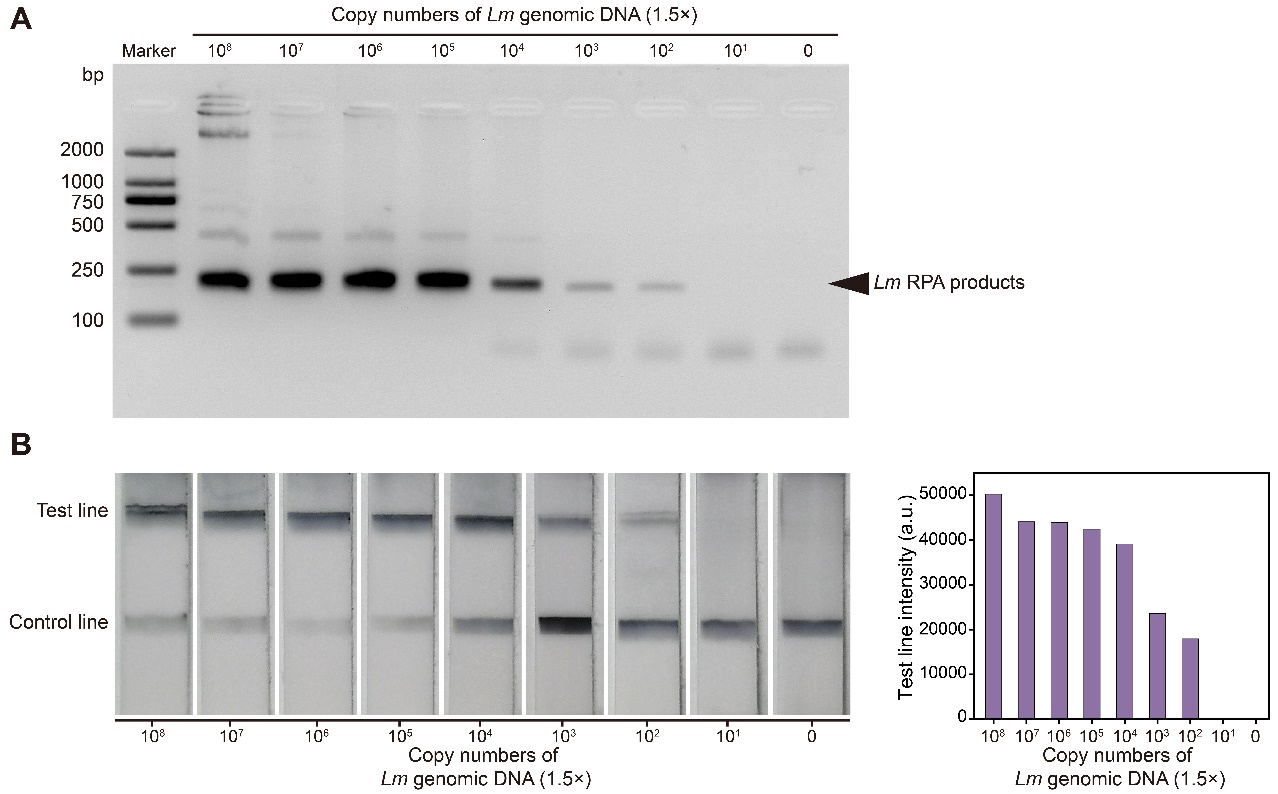

**Figure S3**. **CASLFA detection of *Lm* genomic DNA based on the RPA method.** (A) Agarose gel electrophoresis of *Lm* RPA products. (B) Photographs of strips used for *Lm* detection at various genomic DNA concentrations. The bar chart shows the quantitation of test line intensities of strips shown on the left. *Lm*, *Listeria monocytogenes*.

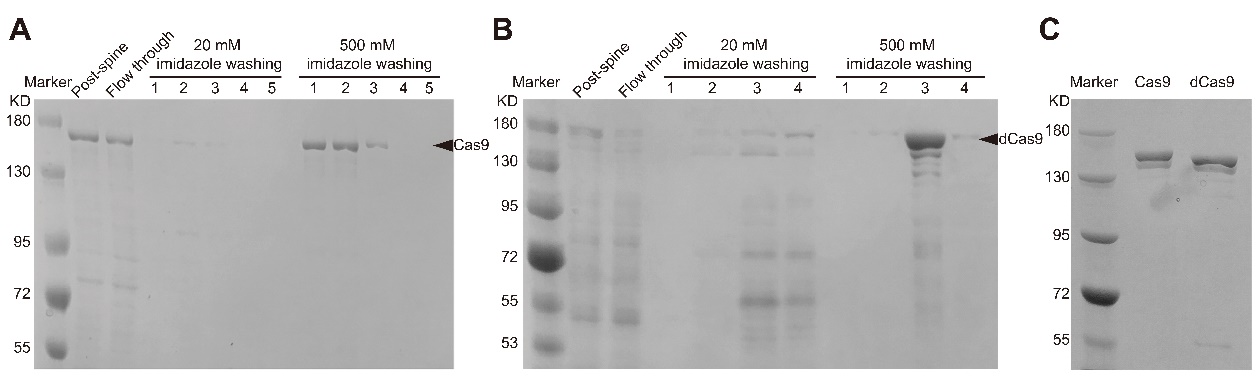

**Figure S4.** **SDS-PAGE analysis of the purified Cas9 and dCas9 proteins.** (A) SDS-PAGE analysis of Cas9 purified with a Ni-NTA column. (B) SDS-PAGE analysis of dCas9 purified with a Ni-NTA column. (C) SDS-PAGE analysis of purified Cas9 and dCas9. Black triangles indicate the Cas9 and dCas9 proteins.

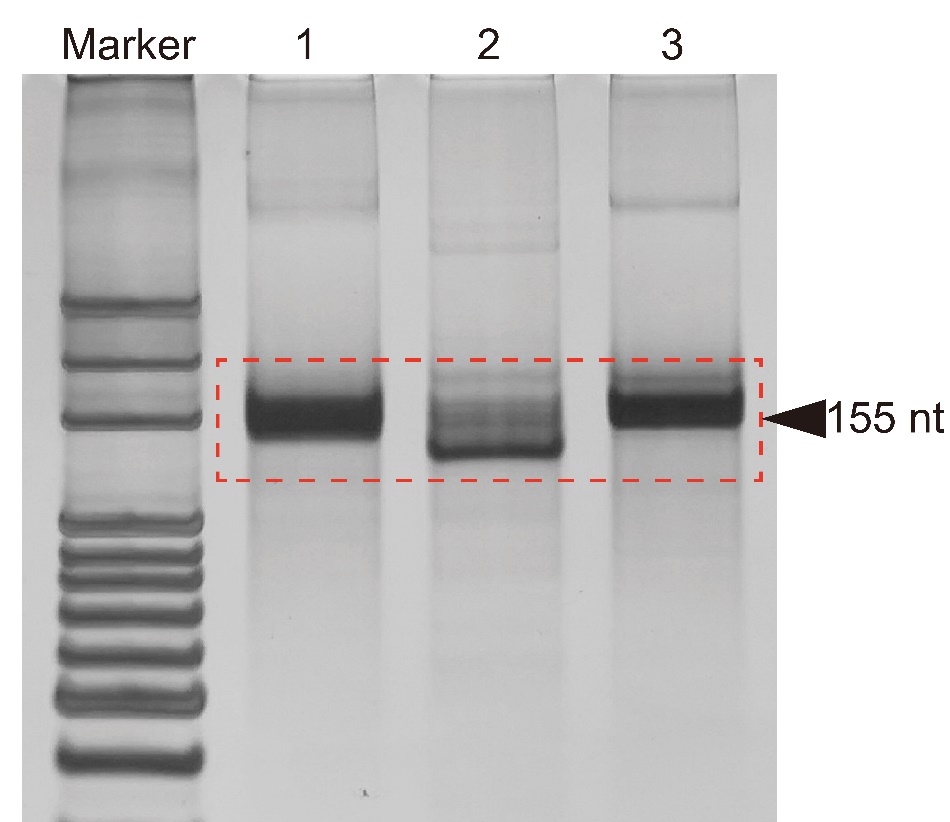

**Figure S5.** **PAGE analysis of the synthesized sgRNAs.** Lane 1: sgRNA used to detect *Listeria monocytogenes*; Lane 2: sgRNA used to detect transgenic rice; Lane 3: sgRNA used to detect African swine fever virus. The bands in the red dashed box are the synthesized sgRNAs with a length of 155 nt.

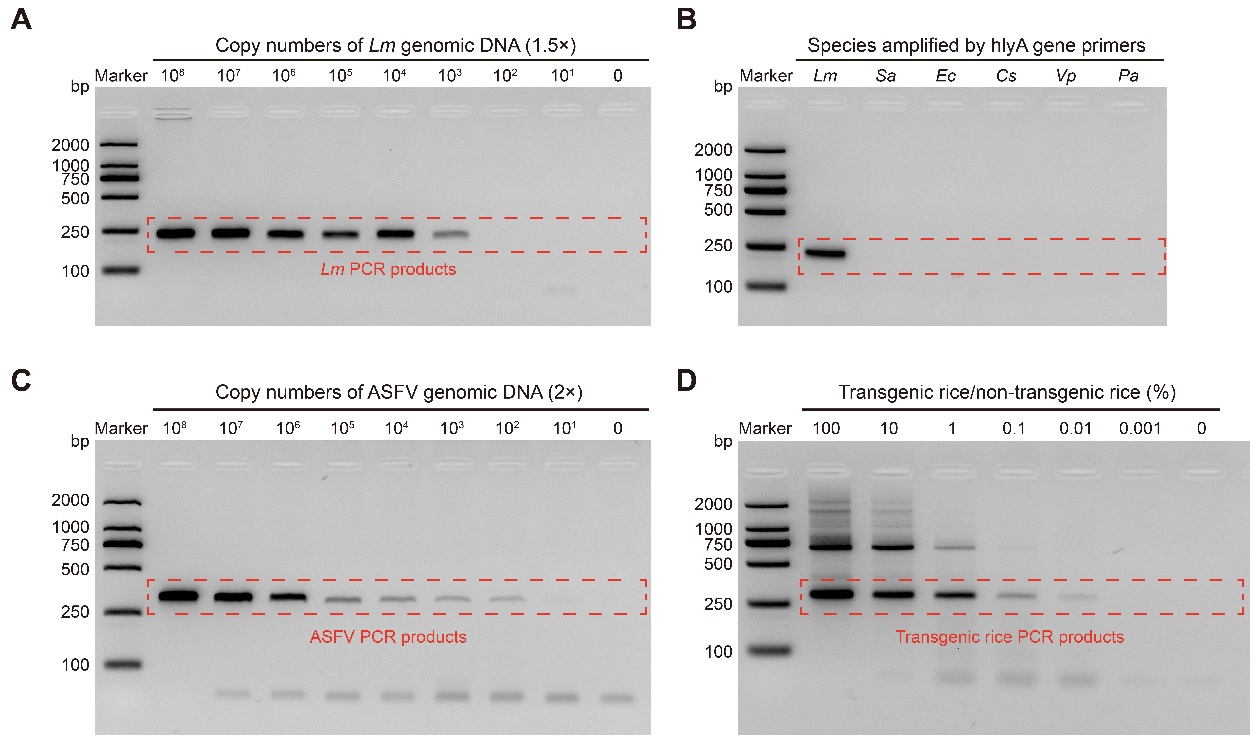

**Figure S6.** **Agarose gel electrophoresis of PCR products.** (A) Agarose gel electrophoresis of *Lm* PCR products. (B) Evaluation of the specificity of amplification using Lm primers by agarose gel electrophoresis. *Lm*, *Listeria monocytogenes*; *Sa*, *Staphylococcus aureus*; *Ec*, *Escherichia coli*; *Cs*, *Cronobacter sakazakii*; *Vp*, *Vibrio parahaemolyticus*; *Pa*, *Pseudomonas aeruginosa*. (C) Agarose gel electrophoresis of African swine fever PCR products. (D) Agarose gel electrophoresis of transgenic rice PCR products.

**Table S1: Nucleic acids sequences used in this work.**

| **Nucleic acids ID** | **Sequences (5’-3’)** |
| --- | --- |
| AuNP-DNA probe 1 | AAAAAAAAAATTTTTCATTGCTAGAGCAGGAGGATGTTGCCGTCCTCCTT |
| AuNP-DNA probe 2 | AAAAAAAAAATTTTTCATTGCTAGAGCAGGGGTATTCCACTTGCA |
| Control line probe | biotin-CCTGCTCTAGCAATG |
| EGFP PCR primer1-F | ATGGTGAGCAAGGGCGAG |
| EGFP PCR primer1-R | TTACTTGTACAGCTCGTCCATGC |
| EGFP PCR primer2-F | biotin-TCCAGGAGCGCACCATCTTC |
| EGFP PCR primer2-F (no biotin) | TCCAGGAGCGCACCATCTTC |
| EGFP PCR primer2-R | TGCCGTTCTTCTGCTTGTCG |
| EGFP sgDNA-F (normal) | CCTCTAATACGACTCACTATAGGAAGGAGGACGGCAACATCCTGTTTAAGAGCTATGCTGGAAACAGCA |
| EGFP sgDNA-R (normal) | AAGAAAAAAAGCACCGACTCGGTGCCACTTTTTCAAGTTGATAACGGACTAGCCTTATTTAAACTTGCTATGCTGTTTCCAGCATAGCTC |
| EGFP sgDNA-F | CCTCTAATACGACTCACTATAGGAAGGAGGACGGCAACATCCTGTTTAAGAGCTATGCTGGAAAAAAGAAAAATGCAAGTGGAATACCAAAAAGA |
| Universal sgDNA-R | AAGAAAAAAAGCACCGACTCGGTGCCACTTTTTCAAGTTGATAACGGACTAGCCTTATTTAAACTTGCTATGCTGTTTTTTTCTTTTTGGTATTCC |
| *Listeria monocytogenes hlyA* gene PCR-F | biotin-CCGTAAGTGGGAAATCTG |
| *Listeria monocytogenes hlyA* gene PCR-R | TTGTTGTATAGGCAATGGG |
| *Listeria monocytogenes hlyA* gene RPA-F | biotin-GTAAGTGGGAAATCTGTCTCAGGTGATGTAG |
| *Listeria monocytogenes hlyA* gene RPA-R | ACTCCTGGTGTTTCCCGGTTAAAAGTAGCA |
| *Listeria monocytogenes hlyA* gene sgDNA-F | CCTCTAATACGACTCACTATAGGGATGAAGTTCAAATCATCGAGTTTAAGAGCTATGCTGGAAAAAAGAAAAATGCAAGTGGAATACCAAAAAGA |
| Transgenic rice 35S promotor-PCR-F | biotin-CCTCCTCGGATTCCATTG |
| Transgenic rice 35S promotor-PCR-R | GGATTGTGCGTCATCCCT |
| Transgenic rice 35S promotor sgDNA-F | CCTCTAATACGACTCACTATAGGAAATGCCATCATTGCGATAAGTTTAAGAGCTATGCTGGAAAAAAGAAAAATGCAAGTGGAATACCAAAAAGA |
| African swine fever p72 gene PCR-F | biotin-ATGGATACCGAGGGAATAGC |
| African swine fever p72 gene PCR-R | CTTACCGATGAAAATGATAC |
| African swine fever p72 gene sgDNA-F | CCTCTAATACGACTCACTATAGGTGATAGTATTTAGGGGTTTGGTTTAAGAGCTATGCTGGAAAAAAGAAAAATGCAAGTGGAATACCAAAAAGA |
